# The MKK3 module integrates nitrate and light signals to modulate secondary dormancy in *Arabidopsis thaliana*

**DOI:** 10.1101/2024.01.28.577345

**Authors:** Sarah Regnard, Masahiko Otani, Marc Keruzore, Alizée Teinturier, Marc Blondel, Naoto Kawakami, Anne Krapp, Jean Colcombet

## Abstract

Seed dormancy corresponds to a reversible blockage of germination. Primary dormancy is established during seed maturation while secondary dormancy is set up on the dispersed seed, following an exposure to unfavourable factors. Both dormancies are relieved in response to environmental factors, such as light, nitrate and coldness. QTL analyses for preharvest sprouting identified MKK3 kinase in cereals as a player in dormancy control. Here, we showed that MKK3 also plays a role in secondary dormancy in Arabidopsis within a signalling module composed of MAP3K13/14/19/20, MKK3 and clade-C MAPKs. Seeds impaired in this module acquired heat-induced secondary dormancy more rapidly than WT seeds and this dormancy is less sensitive to nitrate, a signal able to release dormancy. We also demonstrated that MPK7 was strongly activated in the seed during dormancy release, especially in response to light and nitrate. This activation was greatly reduced in *map3k13/14/19/20* and *mkk3* mutants. Finally, we showed that the module was not regulated, and apparently did not regulate, the genes controlling ABA/GA hormone balance, one of the crucial mechanisms of seed dormancy control. Overall, our work identified a whole new MAPK module controlling seed germination and enlarged the panel of functions of the MKK3-related modules in plants.

## Introduction

A seed is a dispersive organ produced by plants after fertilisation. The plantlet embryo, embedded in the seed, can remain functional—although asleep—for long periods, even in harsh environmental conditions. Its genetic programme, shaped by the species’ evolution in interaction with its environment, allows its germination at the appropriate time to maximise the probability of producing progeny. A critical physiological determinant of seed behaviour is its dormancy (i.e., its ability to block germination even when the environmental conditions are compatible with germination and plant development) and how it reacts to environmental inputs. Therefore, dormancy can be considered as a seed mechanism integrating environmental conditions that will drive the decision to germinate or not. Primary dormancy is acquired during the seed maturation on the mother plant (Baskin and Baskin, 2004). It is the highest when seeds have just been released and gradually decreases with time. Once they reach the ground, and depending on the species’ lifestyle, seeds face a succession of favourable and unfavourable germination periods throughout the year, which requires the repetitive establishment of *de novo* dormancy and its breaking, this process being referred to as ‘dormancy cycling’ (Footitt et al., 2011). This secondary dormancy is acquired by dispersed seeds, which cannot germinate when the conditions are unfavourable. All aspects of seed dormancy, including its acquisition, maintenance, and breaking, are tightly modulated by environmental cues.

Dormancy is thought to be primarily controlled by a dynamic hormonal balance (Shu et al., 2016; Tuan et al., 2018). Abscisic acid (ABA) is a major inducer of seed dormancy, whereas gibberellins (GA) promote germination. Plants impaired in ABA/GA biogenesis or signalling pathways are affected in their ability to control dormancy and germination. After seed dispersal or harvest, the ABA catabolism in imbibed seeds contributes to the release of dormancy and allows the promotion of germination by GA. Many external cues modulate this balance. Seeds perceive and respond to changes in environmental conditions, such as temperature, light, storage periods (after-ripening), moisture content, and nitrate (NO_3_ ). External NO_3_ promotes seed germination at low concentrations in many plant species, acting as both a nutrient and signal (Alboresi et al., 2005). Furthermore, exogenous NO_3_ breaks the primary dormancy efficiently and promotes the completion of seed germination by enhancing ABA catabolism and inhibiting ABA synthesis. In Arabidopsis, the NIN-Like Protein 8 (NLP8) transcription factor is a major regulator of NO_3_ -regulated dormancy, promoting NO_3_ -dependent germination by upregulating ABA catabolism genes that include *CYP707A1/2* (Yan et al., 2016).

In recent years, several reports pointed to the MKK3 kinase as an essential actor promoting seed germination. Notably, MKK3 orthologous genes have been identified in wheat, barley, and rice as the underlying molecular traits for QTLs controlling pre-harvest sprouting (PHS) (Mao et al., 2020; Nakamura et al., 2016; Torada et al., 2016). Arabidopsis seeds impaired in *MKK3* also displayed defects in germination and ABA sensitivity (Danquah et al., 2015). MKK3 belongs to the MAP2K (MAPK Kinase) subfamily which is found in plants and other eukaryotes. Together with MAP3K (MAP2K Kinase) and MAPK (Mitogen-Activated Protein Kinase) subfamilies, it forms intracellular phosphorylation cascades, referred to as MAPK modules (Jagodzik et al., 2018; Colcombet and Hirt, 2008). Our work and others have suggested that the Arabidopsis MKK3 is robustly activated by clade-III MAP3Ks (MAP3K13/14/15/16/17/18/20/21) and, in turn, activates clade-C MAPKs (MPK1/2/7/14). Such modules were largely involved in abiotic stress signalling. Drought, through the ABA core signalling module, activated a MAP3K17/18-MKK3-MPK7 module in plantlets, and plants impaired in these kinases display reduced tolerance to drought (Danquah et al., 2014; Mitula et al., 2015; Li et al., 2017b; Matsuoka et al., 2015; Zhou et al., 2021). Unexpectedly, the module’s activation was tightly controlled by the transcription-dependent MAP3K17/18 production, which, without stimulation, did not occur (Danquah et al., 2015; Boudsocq et al., 2015). More recently, the MKK3 module was found to be activated in response to insect feeding, wounding, and jasmonic acid (Sözen et al., 2020). Again, the module’s activation depended on the strong upregulation of several clade-III *MAP3K* genes. Last, we have recently showed that NO_3_ (nitrate) activates an MKK3 module through the NIN-Like Protein 7 (NLP7)-dependent upregulation of *MAP3K13/14* genes (Sözen et al 2020; Schenk, Chardin, Krapp, and Colcombet, unpublished). In the present work, we further investigate the functioning of the MKK3 module in the control of dormancy using the model plant Arabidopsis.

## Results

### Seeds impaired in *MKK3* establish a faster secondary dormancy

To test whether the MKK3 module plays a role in the acquisition of secondary dormancy, Col-0 and *mkk3-1* seeds produced in low NO_3_ ^-^ conditions were sown on agar plates containing 5 mM KCl and incubated at 30 °C in the dark. Even after 15 days, thermo-inhibition prevented seeds from germinating (*n* > 10). Seeds were then transferred into permissive conditions (16 h photoperiod; 22 °C) for seven days to measure their germination ability. Upon this treatment, Col-0 seeds progressively acquired dormancy, with a 50% decrease in germination after 7–10 days. *mkk3-1* seeds acquired dormancy more rapidly, in less than two days, suggesting that the MKK3 module prevents dormancy acquisition (Figure 1A). Next, we wondered whether the module could function in the release of secondary dormancy. Because NO_3_ was shown to be able to promote germination, we tested whether it could break the heat-induced secondary dormancy. Seeds were first incubated on agar plates containing 5 mM KCl at 30 °C to induce secondary dormancy and then transferred on new plates supplemented with various NO_3_ concentrations and incubated for seven days in permissive conditions (16 h; photoperiod 22 °C). NO_3_ was provided in the form of KNO_3_, and KCl was added to reach a KCl + KNO_3_ concentration of 5 mM, keeping potassium and total anion concentrations constant. As previously shown, seeds were largely dormant when transferred on agar supplemented with only KCl (Figure 1B). *mkk3-1* seeds displayed lower germination on low NO_3_ concentrations (0.05 and 0.5 mM) than Col-0. 5 mM KNO_3_ was able to promote about 100% germination in both genotypes. We found a similar result for *mkk3-2*, and the *mkk3-1* complemented line presented a WT behaviour (Figures S1C and S1D). This result indicates that *mkk3* seeds are less sensitive to NO_3_ ^-^ although not insensitive, suggesting that an MKK3-containing MAPK module could mediate a part of the NO_3_ signalling to promote dormancy breaking.

**Figure 1.**
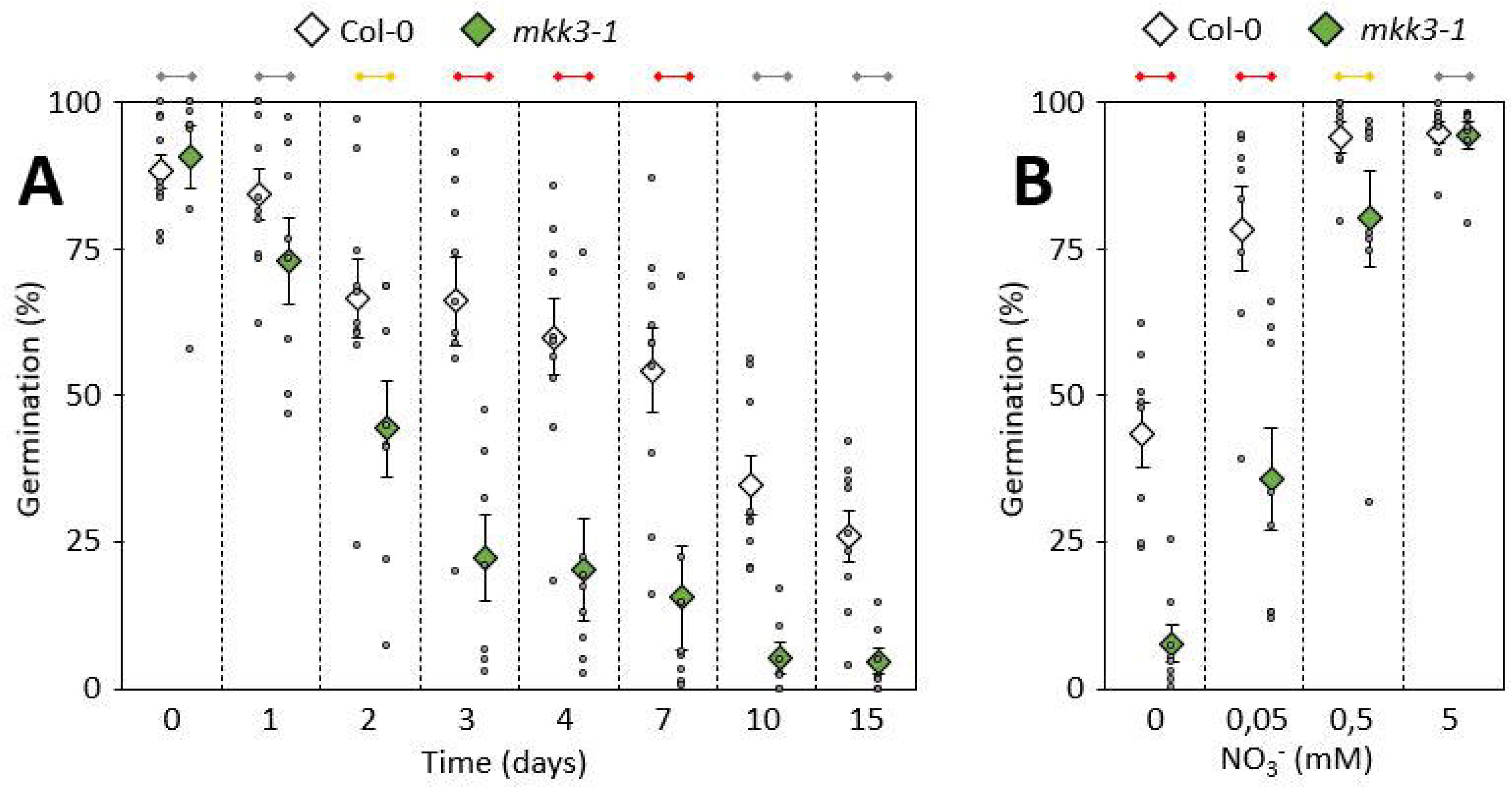
*mkk3-1* seeds have a faster secondary dormancy acquisition and a reduced nitrate-triggered dormancy release. A. Col-0 and *mkk3-1* seeds were imbibed at 30°C in the dark for the indicated time to induce secondary dormancy. Germination ability was then assessed after 7 days in long day conditions. Values are mean ± SE of seven to ten biological replicates from seed batches produced independently. Values for each replicate are also shown. On the top, based on Mann-Whitney test, yellow and red sticks show differences with 1% < α < 5% and α < 1%, respectively, whereas gray sticks show no differences. B. Col-0 and *mkk3-1* seeds were imbibed at 30°C in the dark for 10 days to induce secondary dormancy and transferred on media containing indicated NO3-concentrations. Germination ability was then assessed after 7 days in long day conditions. Values are mean ± SE of height biological replicates from seed batches largely produced independently. Values for each replicate are also shown. On the top, based on Mann-Whitney test, yellow and red sticks show differences with 1% < α < 5% and α < 1%, respectively, whereas gray sticks show no differences.

### *MPK7* is expressed in seeds and activated by NO ***^-^*** in an MKK3-dependent way

MKK3 was shown to activate MAPKs of the clade-C, namely MPK1/2/7/14 (Colcombet et al., 2016; Danquah et al., 2015; Sözen et al., 2020). We evaluated by RT-qPCR analysis which of these clade-C MAPKs were expressed in seeds during the first hours of germination after transfer on 5 mM KCl or 5 mM KNO_3_ (Figure 2). *MPK1* and *MPK7* displayed a higher expression than *MPK2* and *MPK14*. None of the four clade-C MAPK genes seemed to be more sensitive to NO_3_^-^ than to Cl^-^, and all four displayed a decrease in their expression after transfer to germination conditions. HA-tagged MPK1/2/7-locus lines available in the laboratory (Sözen et al., 2020) confirmed that MPK7 was detectable in dry seeds but suggested that MPK1 and MPK2 were not (Figure S2). Seeds impaired in the four clade-C MAPKs displayed an *mkk3*-like phenotype in nitrate-triggered dormancy release, whereas seeds impaired in only *MPK7* (single mutant) had an intermediate phenotype (Figure 3). These genetic data are in good agreement with clade-C MAPKs working downstream of MKK3, suggests a functional redundancy in the modulation of secondary dormancy, and indicate that MPK7 can be used as a proxy to monitor the module’s activation.

**Figure 2.**
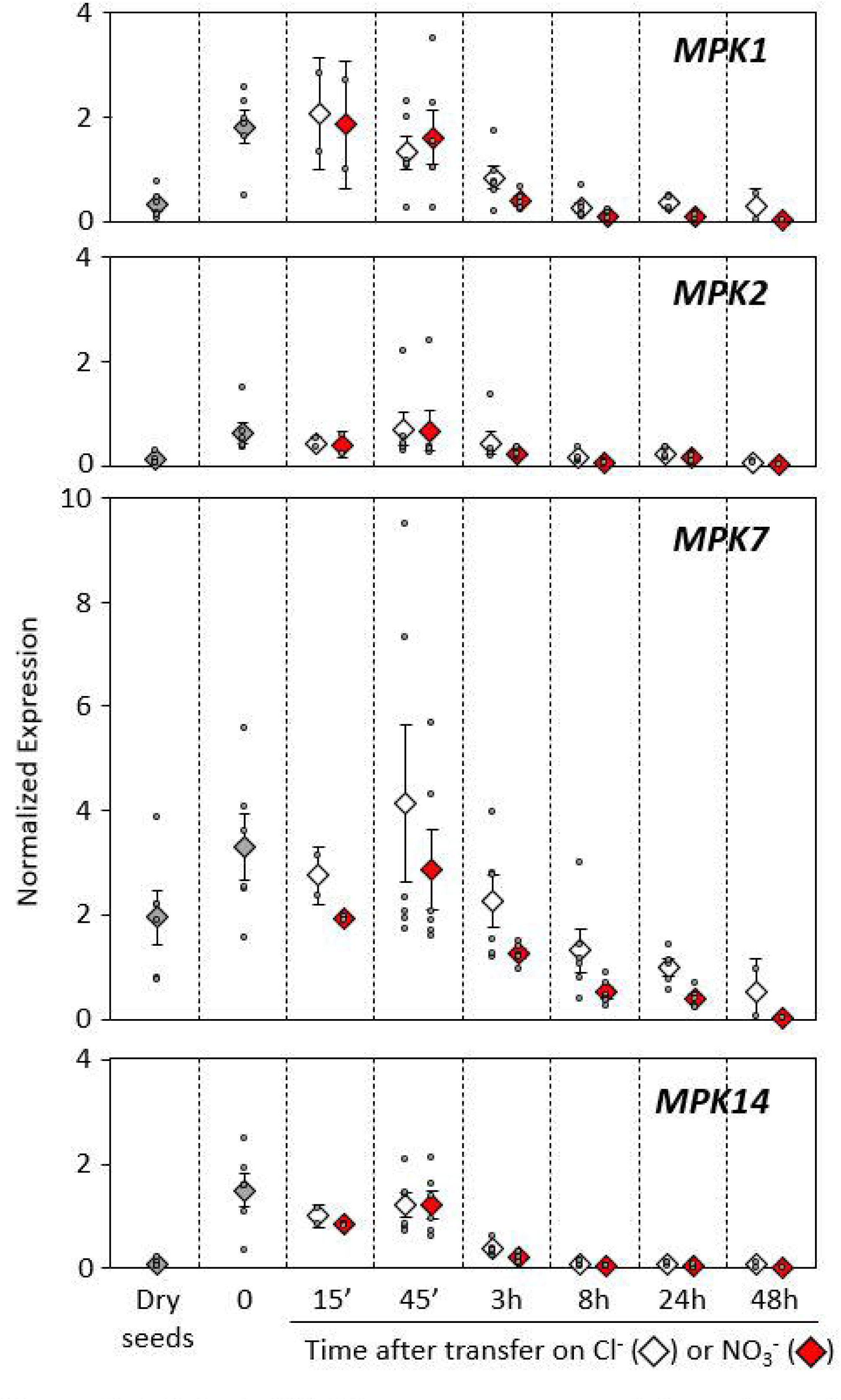
Clade-C *MAPK* genes are expressed in seeds and during secondary dormancy release. RT-qPCR analysis of clade-C MAP3K genes. Transcript levels are expressed relative to ACTIN2 as reference gene. Values are mean ± SE of two to six biological replicates from seed batches produced independently. Values for each replicate are also shown.

**Figure 3.**
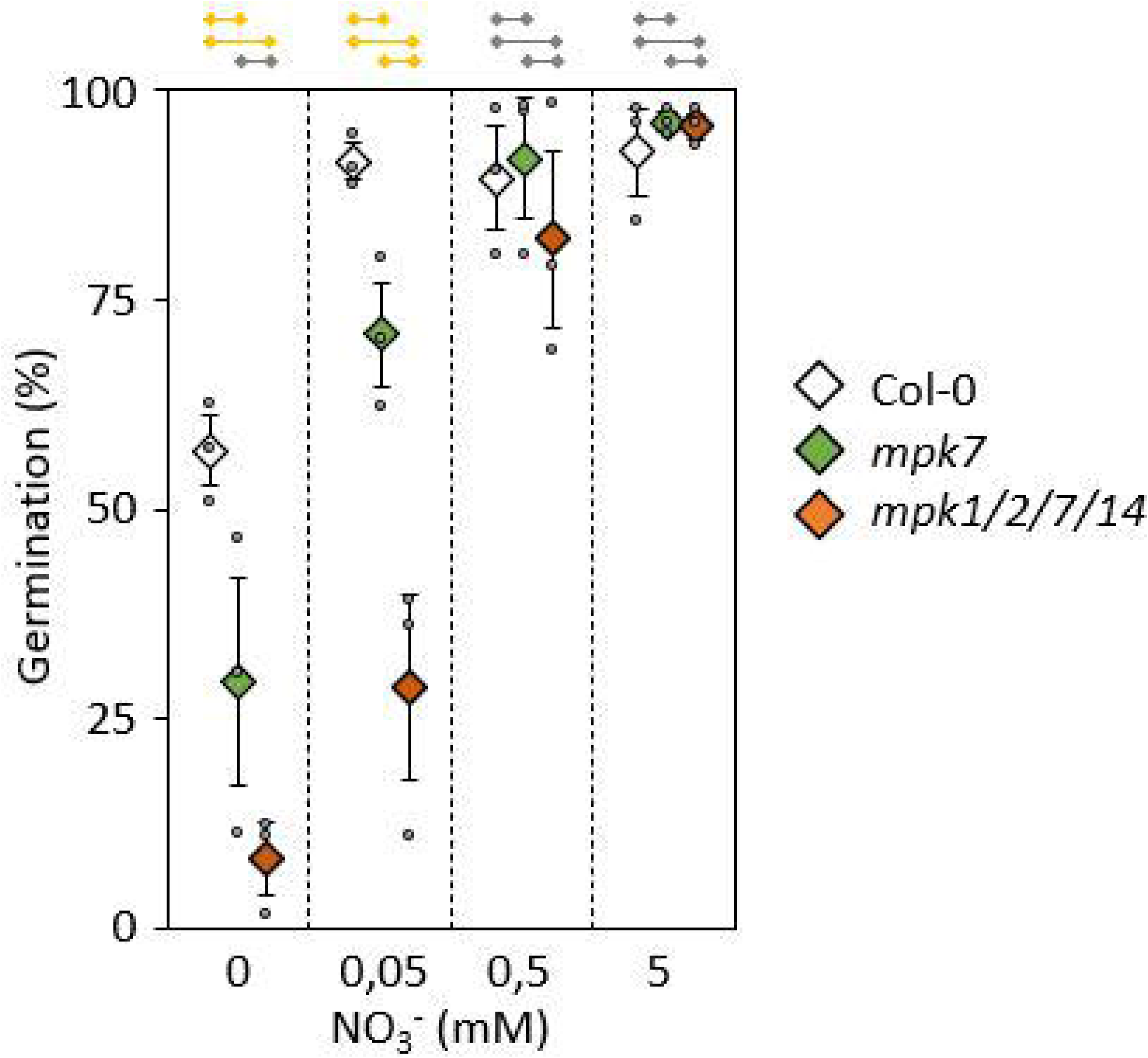
Seeds impaired in C-clade MAPKs have a faster secondary dormancy acquisition and a reduced nitrate-triggered dormancy release. Col-0, *mpk7* and *mpk1/2/7/14* seeds were imbibed at 30°C in the dark for 10 days to induce secondary dormancy and transferred on media containing indicated NO3-concentrations. Germination ability was then assessed after 7 days in long day conditions. Values are mean ± SE of three biological replicates from seed batches largely produced independently. Values for each replicate are also shown. On the top, based on Mann-Whitney test, yellow sticks show differences with α < 5% whereas gray sticks show no differences.

To assess the module activation, MPK7 was immunoprecipitated from seeds using a specific antibody, and its activity was assayed as the ability to phosphorylate the heterologous substrate MYELIN BASIC PROTEIN (MBP) (Figures 4 and S3). MPK7 activity was detectable in dry seeds but not in seeds under heat-induced dormancy. When seeds were transferred to germination conditions (22 °C under light), the activity increased rapidly with an apparent maximum at 8 hours. This increase of MPK7 activity was 3-4 times higher when seeds were transferred on agar supplemented with 5 mM KNO_3_ than agar supplemented with 5 mM KCl (figure S3). In *mkk3-1* seeds, MPK7 activity was not detectable, confirming that MPK7 functions downstream of MKK3. In the *mpk7* background, MPK7 activity was not detectable, confirming the specificity of the antibody raised against MPK7 used in kinase assays and western blots (Figure S4). To consolidate these results, an HA-antibody was used to immunoprecipitate MPK7-HA from plants transformed with an HA-tagged MPK7-locus. MPK7-HA displayed a similar pattern of activity, which was abolished when the *mkk3-1* mutation was introgressed in the line (Figure S5). Overall, these results indicate that MPK7 in seeds is activated by NO_3_ in an MKK3-dependent way. Other activators may be responsible for the high activity background observed when seeds were transferred on KCl.

**Figure 4.**
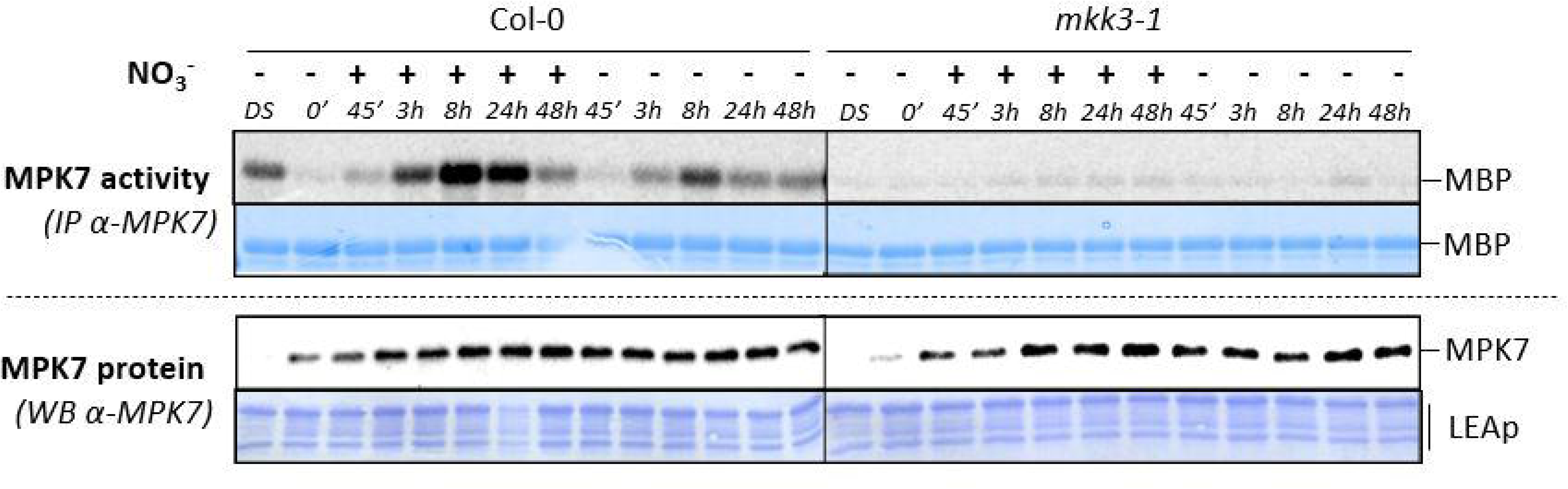
MPK7 activity in dormant seeds transferred on nitrate depends on MKK3. Kinase activity of MPK7 after immunoprecipitation with an anti-MPK7 antibody from Col-0 and mkk3-1 seeds, either dry (DS), after acquisition of secondary dormancy (0’) and after transfer on either 5mM KCl (-) or KNO3. MPK7 amount was monitored by immunoblot using an anti-MPK7 antibody. Equal loading was controlled by Coomassie staining of the membrane. LEAp, Late Embryogenesis Abundant proteins. Results were repeated two to six times depending on the time points, the quantification of these replicates being gathered in figure S3.

### MAP3K13/14/19/20 are necessary for the MKK3-dependent module’s activation

We reported that the activation of MKK3 relies on the transcriptional upregulation of clade-III MAP3Ks (Colcombet et al., 2016; Danquah et al., 2015; Sözen et al., 2020). Using RT-qPCR, we measured the expression of clade-III *MAP3K* genes in Col-0 seeds during dormancy breaking (Figures 5 and S6). Four MAP3K genes were repeatedly upregulated in at least one of the samples. *MAP3K13* and *MAP3K14* were rapidly induced in seeds by NO_3_ ^-^, with a peak at 45 minutes (Figures 5A and 5B), whereas *MAP3K19* and *MAP3K20* displayed a delayed activation, typically at 3 and 8 hours, which was stronger in the presence of 5 mM KNO3 than 5 mM KCl (Figures 5C and 5D). Additionally, *MAP3K13* and *MAP3K19* displayed high expression levels in dry seeds but were not expressed anymore in seeds under heat-induced dormancy (Figures 5A et 5C). These results fit the timing of MPK7 activity in seeds (Figure 4) and suggest that those four MAP3Ks may play a role in the context of seed dormancy regulation. Nevertheless, the patterns of MPK7 activation and *MAP3K* transcriptional regulation suggest a prominent function for MAP3K19/20, whereas the *MAP3K13/14* transcription peak does not coincide with a high MPK7 activation level.

**Figure 5.**
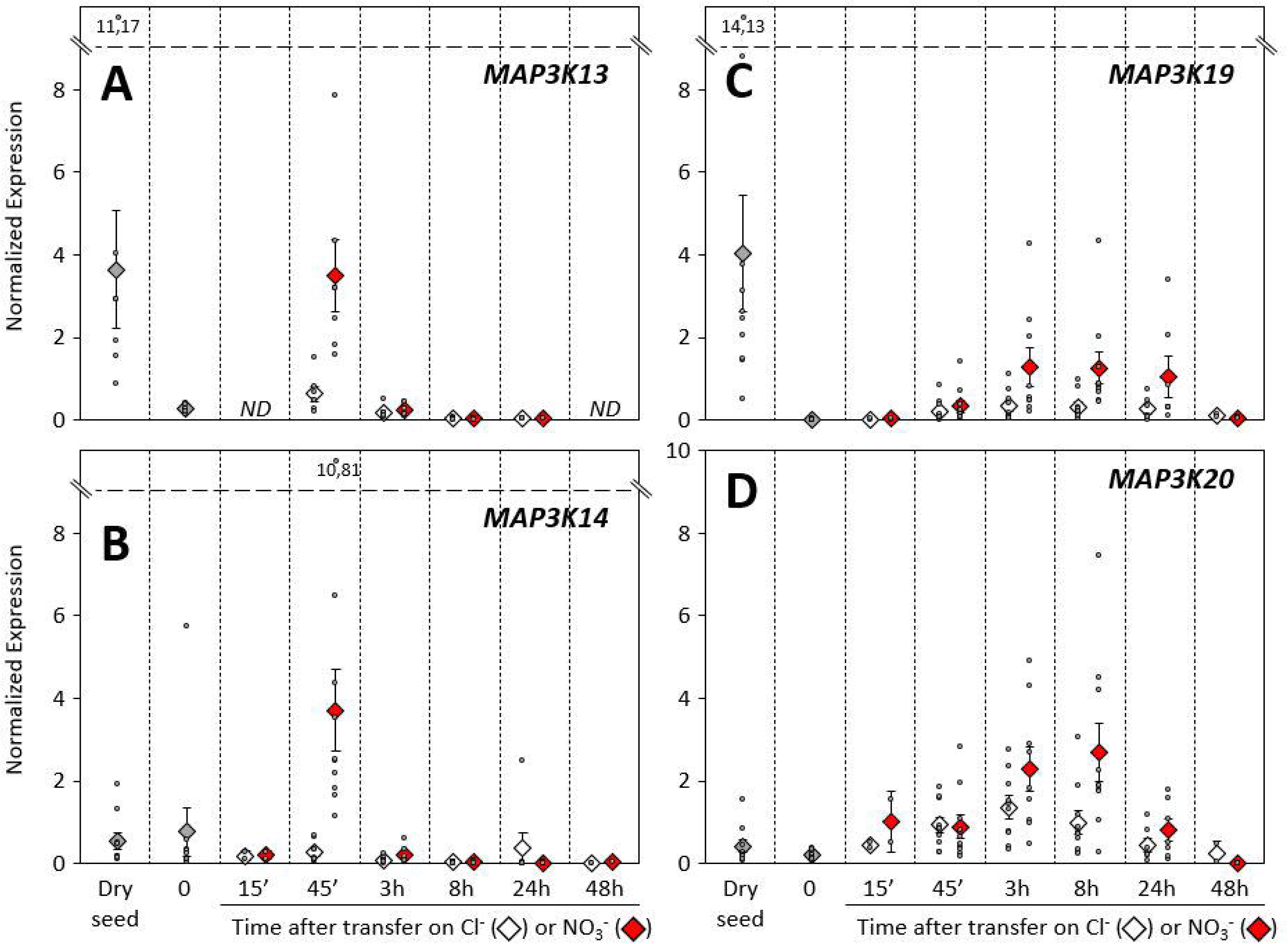
*MAP3K13*, *MAP3K14*, *MAP3K19* and *MAP3K20* are expressed in seeds and during secondary dormancy release. RT-qPCR analysis of *MAP3K13*, *MAP3K14*, *MAP3K19* and *MAP3K20* genes expression. Transcript levels are expressed relative to ACTIN2 as reference gene. Values are mean ± SE of two to 10 biological replicates from seed batches produced independently. Values for each replicate are also shown. ND not determined.

To test the role of these MAP3Ks in seed, we first immunoprecipitated MPK7 from *map3k19-1/20-3* single and double mutants from dormant seeds transferred on KCl or KNO3. MPK7 activity was weakly impaired in *map3k20-3* and strongly in *map3k19-1/20-3*, notably at 8 hours (Figures 6A and S7A). Interestingly, a residual NO_3_ -dependent MPK7 activity was observed in *map3k19-1/20-3* at 45 minutes. Since this residual kinetics fitted the *MAP3K13/14* transcriptional response (figures 5A and 5B), we generated by cross the quadruple mutant *map3k13cr/14cr/19-1/20-3* and showed that the MPK7 activity was impaired throughout the time-course (Figure 6B and S7B). Consistently, *map3k13cr/14cr/19-1/20-3* seeds displayed a stronger NO_3_ ^-^ insensitivity than *map3k13cr/14cr* and *map3k19-1/20-3* seeds (Figure 7). All together, these data suggested that MAP3K13/14/19/20 are the only upstream activators of the nitrate-activated MKK3-MPK1/2/7/14 module involved in the secondary dormancy breaking. To test, whether the MAP3K expression is sufficient to reduce dormancy, we generated two lines constitutively expressing a MYC-tagged *MAP3K19*. They both displayed a higher MPK7 activity (Figures 8A and S8) and did not acquire any secondary dormancy (Figure 8B).

**Figure 6.**
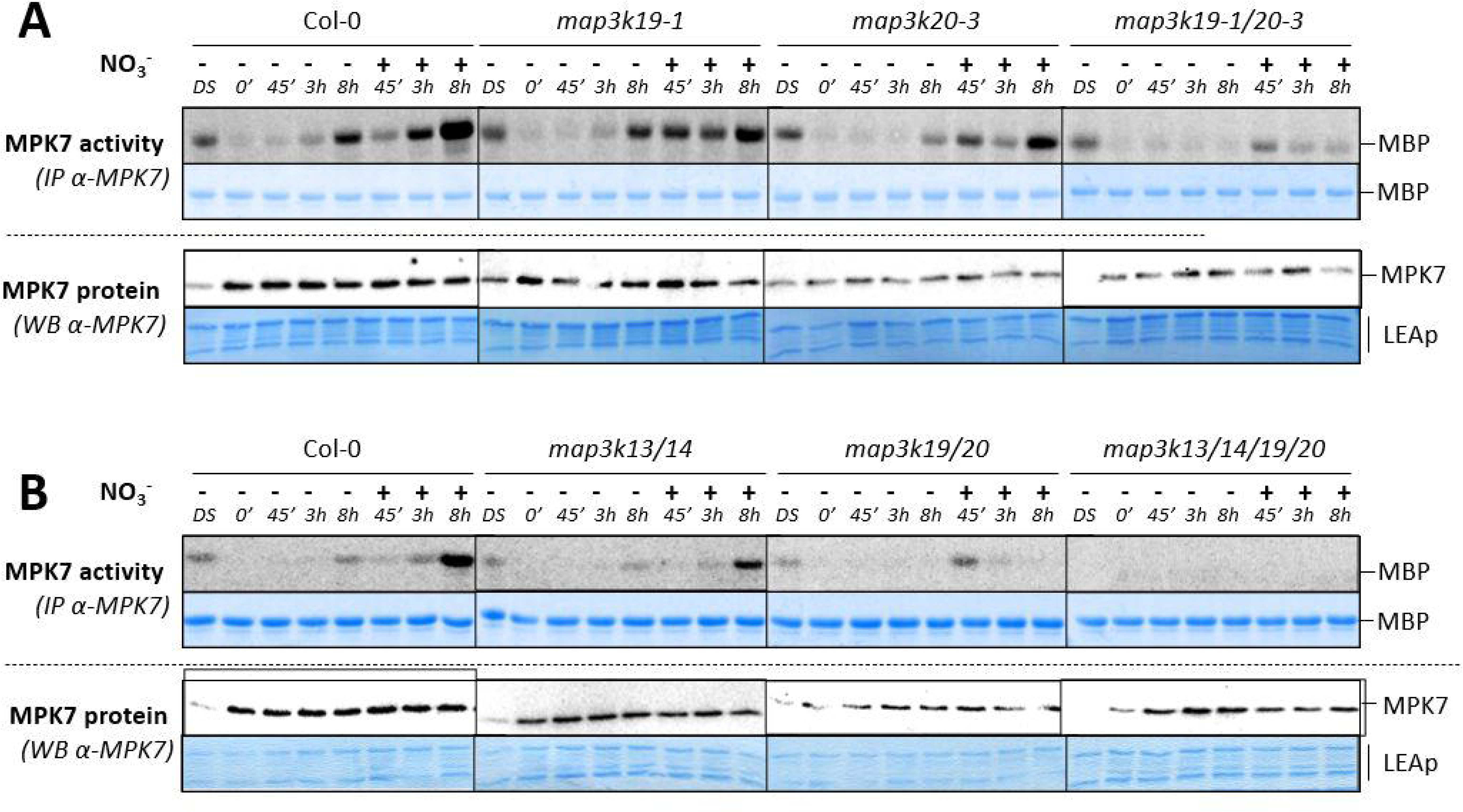
MPK7 activity in dormant seeds transferred on nitrate depends on MAP3K13/14/19/20. Kinase activity of MPK7 after immunoprecipitation with an anti-MPK7 antibody from indicated background, either dry (DS), after acquisition of secondary dormancy (0’) and after transfer on either 5mM KCl or KNO3. MPK7 amount was monitored by immunoblot using an anti-MPK7 antibody. Equal loading was controlled by Coomassie staining of the membrane. LEAp, Late Embryogenesis Abundant proteins. Results were repeated 2-6 times depending of the time point and genotype for A and 3 times for B, the quantification of these replicates being gathered in figures S7A and B.

**Figure 7.**
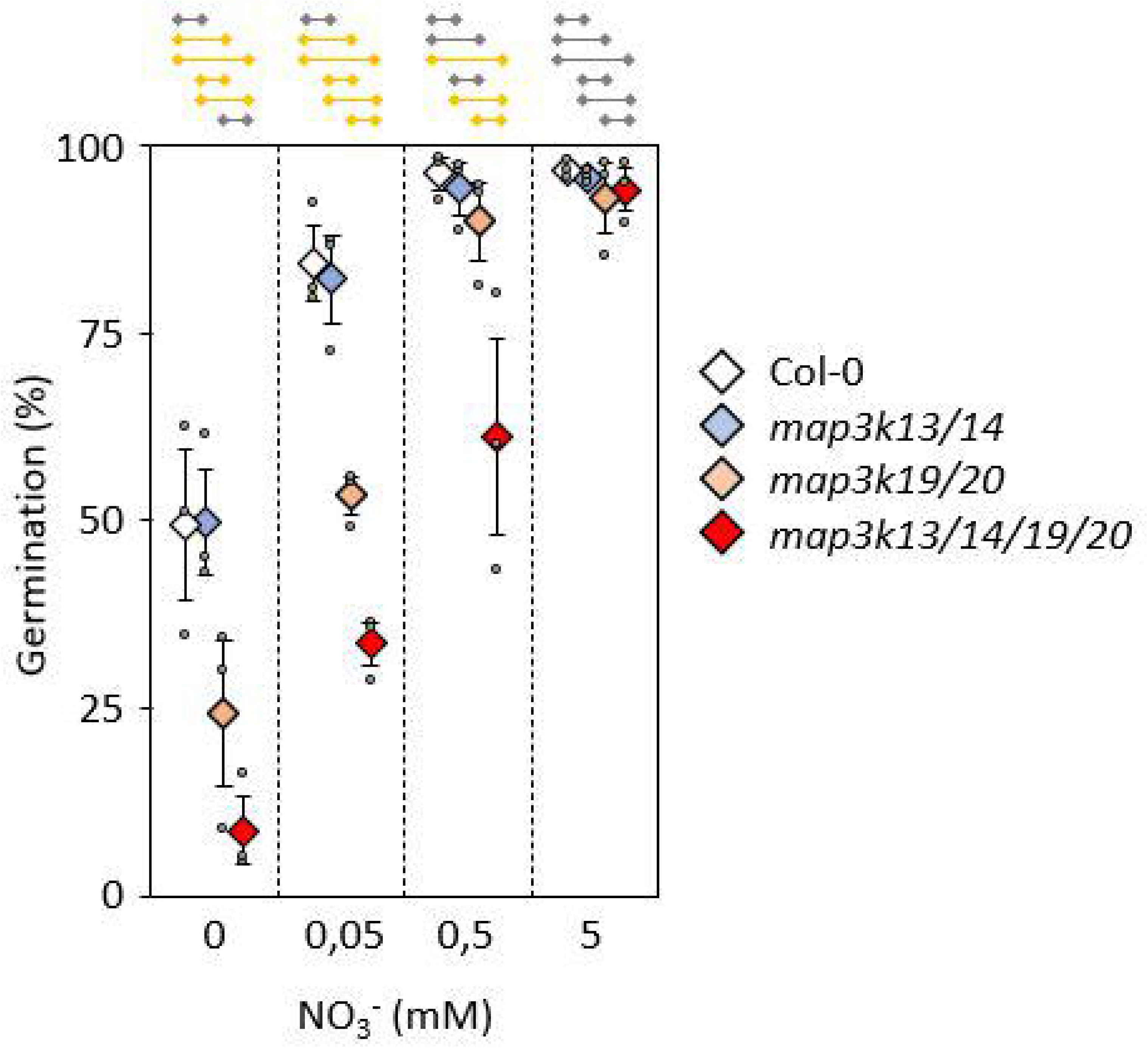
*map3k13/14/19/20* seeds have a reduced nitrate-triggered dormancy release. Col-0, *map3k13CR/14CR, map3k19-1/20-3* and *map3k13CR/14CR/19-1/20-3* seeds were imbibed at 30°C in the dark for 10 days to induce secondary dormancy and transferred on medium containing indicated NO3 concentration. Germination ability was assessed after 7 days in long day conditions. Values are mean ± SE of three biological replicates from seed batches produced independently. Values for each replicate are also shown. On the top, based on Mann-Whitney test, yellow sticks show differences with α < 5% whereas gray sticks show no differences.

**Figure 8.**
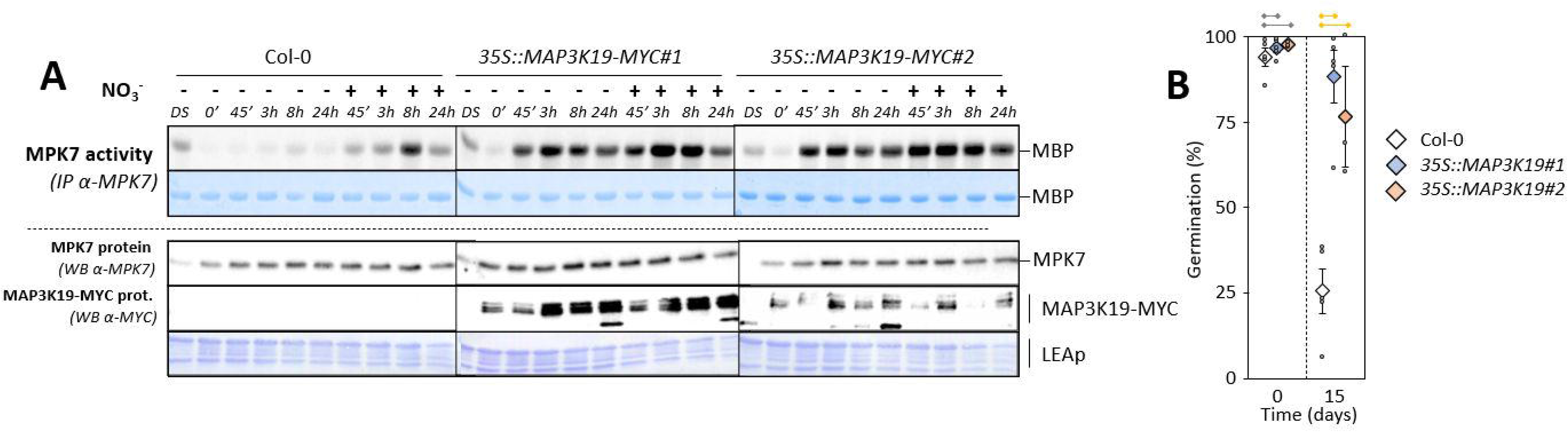
Constitutive expression of *MAP3K19* triggers a strong MKK3-dependent MPK7 activation and reduces the acquisition of secondary dormancy. A. Kinase activity of MPK7 after immunoprecipitation with an anti-MPK7 antibody from indicated background, either dry (DS), after acquisition of secondary dormancy (0’) and after transfer on either 5mM KCl (-) or KNO3 (+). MPK7 amount was monitored by immunoblot using an anti-MPK7 antibody. Equal loading was controlled by Coomassie staining of the membrane. LEAp, Late Embryogenesis Abundant proteins. Results were repeated two to three times depending on the time points, the quantification of these replicates being gathered in figure S8. B. Seeds from indicated background were imbibed at 30°C in the dark for the indicated time to induce secondary dormancy. Germination ability was assessed after 7 days in long day conditions. Values are mean ± SE of 3-4 biological replicates from seed batches produced independently. Values for each replicate are also shown.

### MPK7 activation by nitrate depends of NLP transcription factors

Nitrate has been shown to trigger *MAP3K13/14* expression through NLP transcription factors (Marchive et al., 2013; Yan et al., 2016). To test whether NLPs are also involved in the MKK3 module activation during the breaking of secondary dormancy, we immunoprecipitated MPK7 from dormant seeds impaired in one or several *NLP* genes. Surprisingly, MPK7 activity was not reproducibly reduced in *nlp8* seed sets produced independently (figure S9), whereas NLP8 has been shown to be a master regulator of the nitrate regulation of primary dormancy (Yan et al., 2016). This suggest that other NLPs could act redundantly of NLP8, depending of seed sets. In line, when mutations in other NLPs where combined, MPK7 activity decreased robustly, indicating that NLPs play a role in the *MAP3K* transcriptional regulation by nitrate and therefore in the activation of the module (figures 9 and S10).

**Figure 9.**
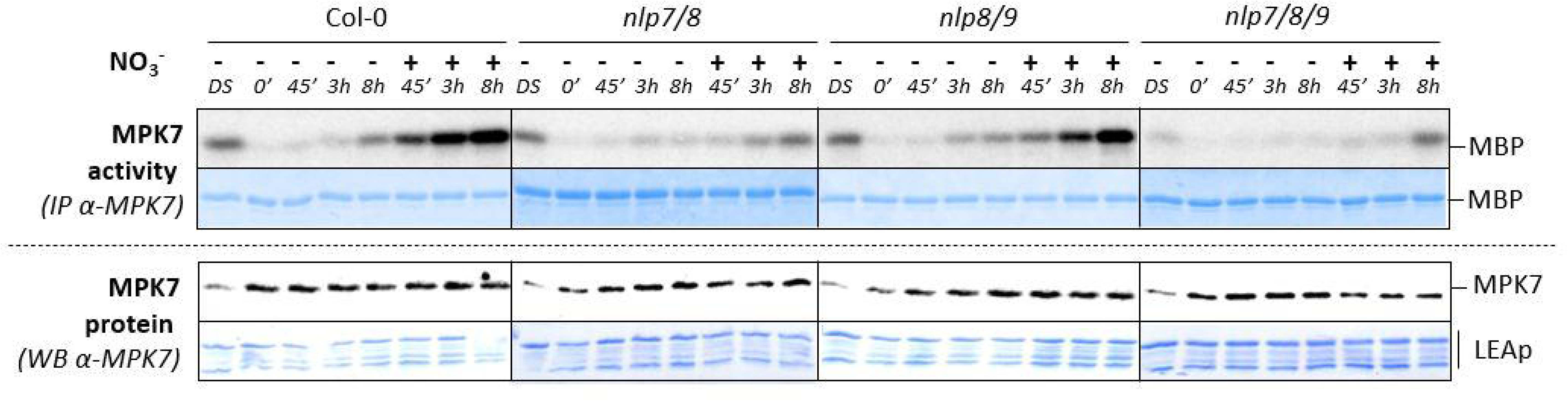
MPK7 activity in dormant seeds transferred on nitrate depends on NLPs. Kinase activity of MPK7 after immunoprecipitation with an anti-MPK7 antibody from indicated background, either dry (DS), after acquisition of secondary dormancy (0’) and after transfer on either 5mM KCl or KNO3. MPK7 amount was monitored by immunoblot using an anti-MPK7 antibody. Equal loading was controlled by Coomassie staining of the membrane. LEAp, Late Embryogenesis Abundant proteins. Results were repeated 3 times, the quantification of these replicates being gathered in figure S10.

### Light activates MPK7 in an MKK3-dependent way and primes *MAP3K19/20* expression

Once treated with heat, seeds were transferred in light conditions, so we wondered whether light might be an activator of the MKK3 module and responsible for the high MPK7 activity background in KCl conditions (see for example, figure 4). We repeated the experiment in dark conditions, manipulating filters carrying dormant seeds under green illumination. In these conditions, we barely detected a background MPK7 activity (Figures 10A and S11). Surprisingly, we did not observe a strong NO_3_ -induced MPK7 activation either, suggesting that the NO_3_ ability to activate MPK7 depends on light. Consistently, RT-qPCR analysis revealed that the transcriptional regulation of *MAP3K19* and *MAP3K20* was strongly reduced in the dark, no matter the anionic condition (Figure 11). On the contrary, the NO_3_ - induced *MAP3K13* and *MAP3K14* expressions were unaffected by light conditions. Other clade-III *MAP3K* genes were not upregulated in these conditions (Figure S12). These results suggest that the *MAP3K19/20* transcriptional regulation acts as a conditional switch for light and NO_3_ in the activation of MPK7 and that the MKK3 module might contribute to the well-known light-triggered germination. Coherently, MPK7 was far less activated by NO_3_ or light in *phyA/B* seeds, impaired in the corresponding phytochrome receptors involved in the red/far-red regulation of germination (Figure 10B).

**Figure 10.**
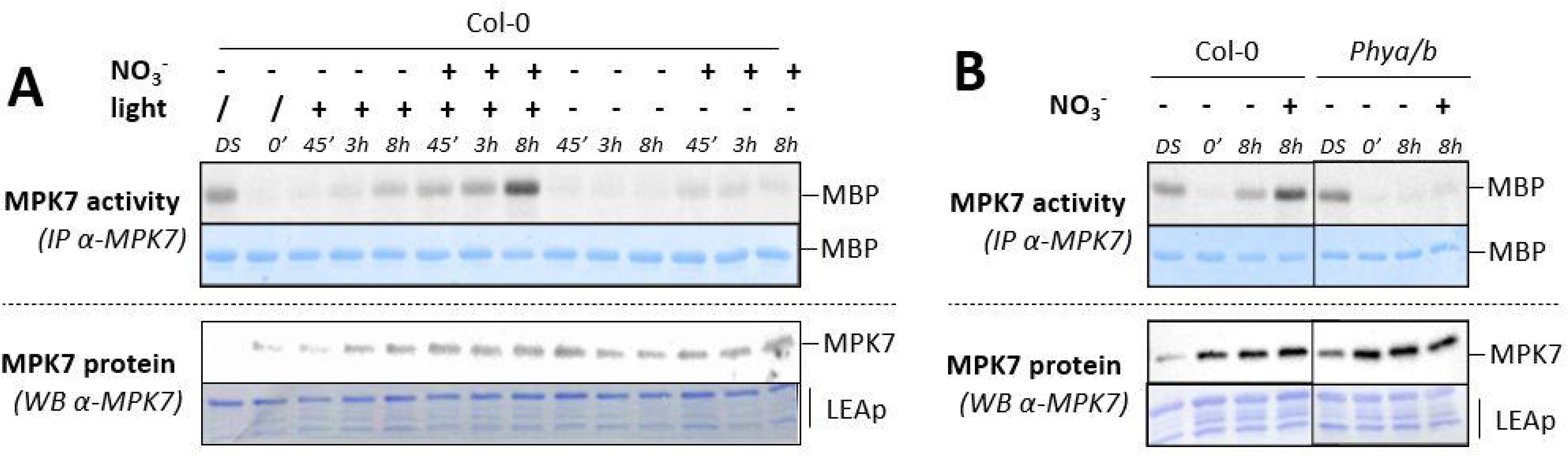
MPK7 activity in dormant seeds is also triggered by light. Kinase activity of MPK7 after immunoprecipitation with an anti-MPK7 antibody from Col-0 (A and B) or *phya/b* (B) seeds, either dry (DS), after acquisition of secondary dormancy (0’) and after transfer on either 5mM KCl or KNO3 with and without whit light. MPK7 amount was monitored by immunoblot using an anti-MPK7 antibody. Equal loading was controlled by Coomassie staining of the membrane. LEAp, Late Embryogenesis Abundant proteins. Results from A were repeated two to three times depending on the time points, the quantification of these replicates being gathered in figure S7. Results from A

**Figure 11.**
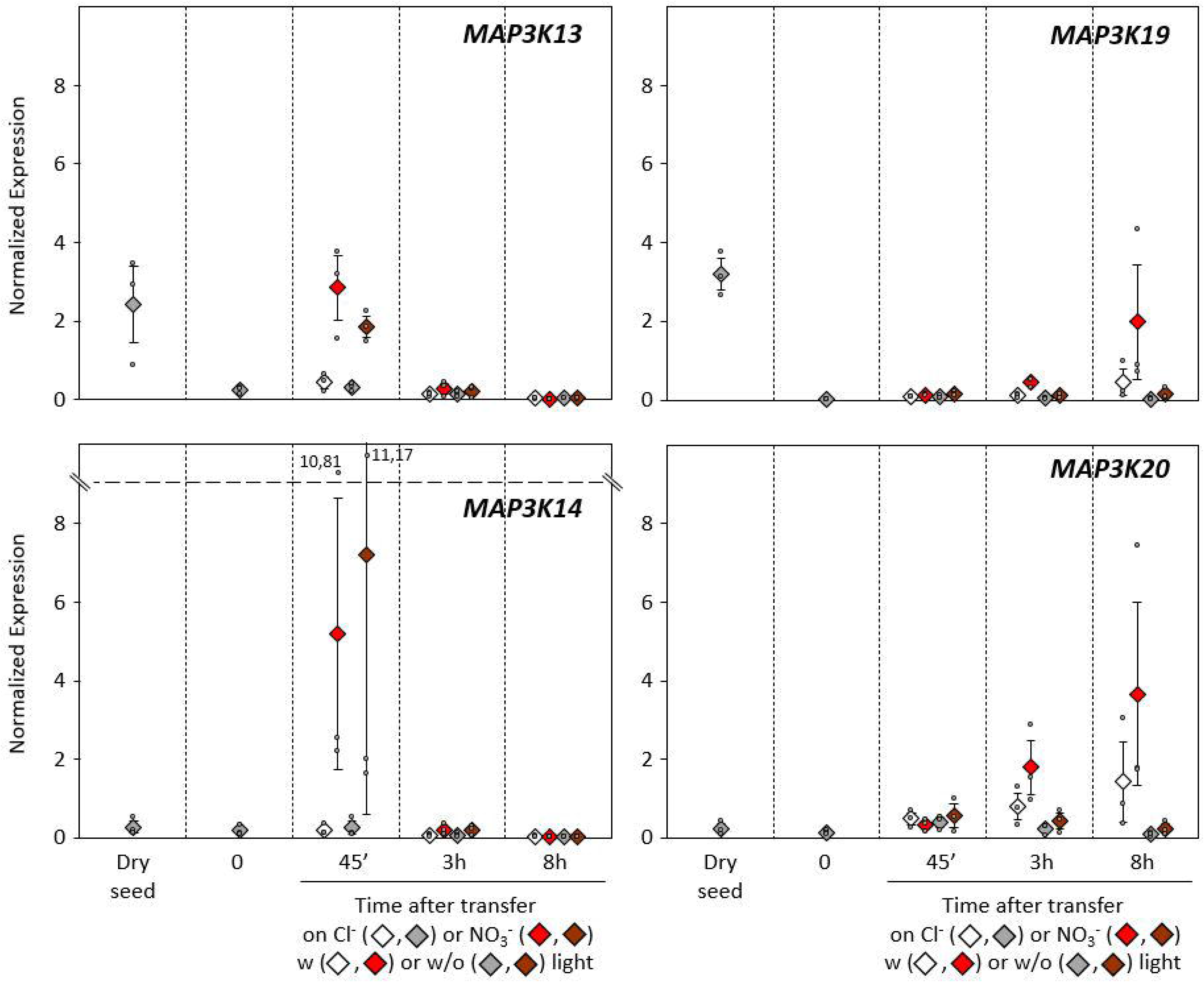
*MAP3K19* and *MAP3K20*, but not *MAP3K13* and *MAP3K14*, are differentially regulated by light. RT-qPCR analysis of *MAP3K13*, *MAP3K14*, *MAP3K19* and *MAP3K20* genes expression. Transcript levels are expressed relative to ACTIN2 as reference gene. Values are mean ± SE of three biological replicates from seed batches produced independently. Values for each replicate are also shown. ND, not determined.

### The MKK3 module activation does not depend on the ABA/GA balance

The ABA/GA hormonal balance is one of the main physiological mechanisms controlling the seed’s decision to germinate. Since the MKK3 module promotes germination, we first wondered whether the NO_3_ - and light-triggered activation of the MKK3 module depended on GA synthesis or signalling. To test this possibility, we transferred dormant seeds on agar media containing either KCl or KNO3, combined or not with 10 µM paclobutrazol (PCZ), a potent blocker of GA biosynthesis. In these conditions, we did not measure any variation in MPK7 activity (Figures 12A and S13A). We also confirmed that an active GA, GA_3_, could not directly activate the module (Figures 12B and S13B).

**Figure 12.**
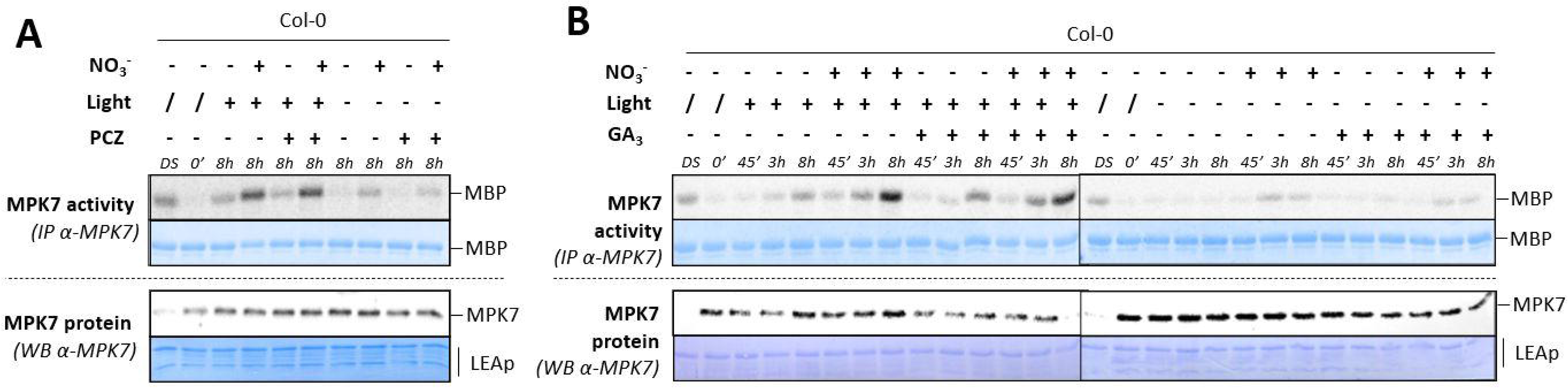
MPK7 activity in dormant seeds is not modulated by Gas. A and B. Kinase activity of MPK7 after immunoprecipitation with an anti-MPK7 antibody from Col-0 seeds, either dry (DS), after acquisition of secondary dormancy (0’) and after transfer on either 5mM KCl or KNO3 with and without white light with and without Paclobutrazol 10µM (A) or with and without GA3 50µM (B). MPK7 amount was monitored by immunoblot using an anti-MPK7 antibody. Equal loading was controlled by Coomassie staining of the membrane. LEAp, Late Embryogenesis Abundant proteins. Results were repeated two to three times depending on the time points, the quantification of these replicates being gathered in figure S13.

We then tested whether ABA could modulate MPK7 activity in seeds. Dormant seeds were transferred on agar containing 5 mM KCl or KNO3, with or without ABA, for 8 hours, and MPK7 activity was assayed. ABA did not affect the NO_3_ -induced MPK7 activation (Figure 13). This result was rather surprising since we previously reported that ABA could activate the module in plantlets through the transcriptional regulation of *MAP3K17/18* (Danquah et al., 2015). When ABA was added to the media, no matter the anionic conditions, seeds did not express *MAP3K17/18* (Figure S14). At the same time, *MAP3K13/14/19/20* displayed unchanged transcriptional regulations (Figure S14). This result indicates that the ABA-dependent activation of the MKK3 module is tissue-specific and that the ABA-dependent regulation of seed germination does not require the MKK3 module in our conditions.

**Figure 13.**
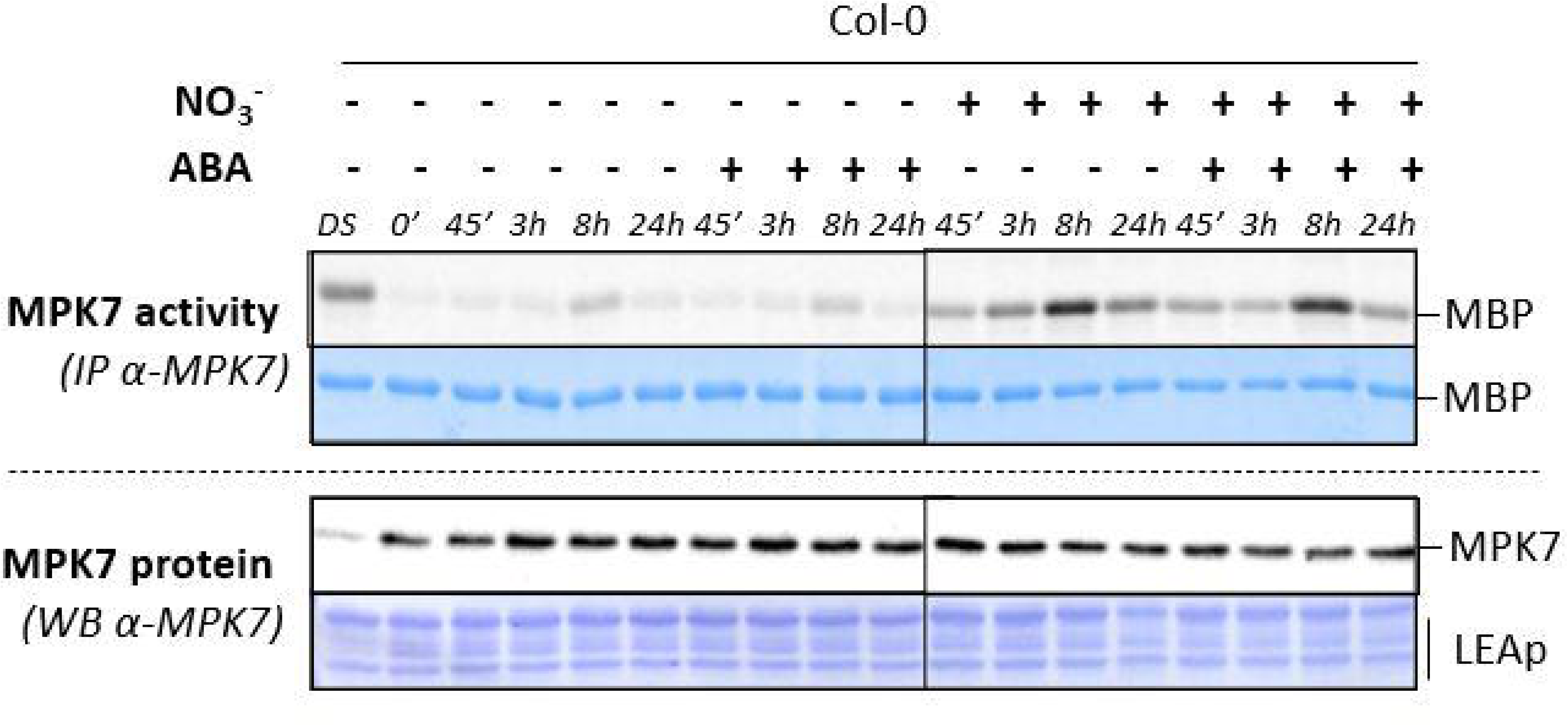
MPK7 activity in dormant seeds is not modulated by ABA. Kinase activity of MPK7 after immunoprecipitation with an anti-MPK7 antibody from Col-0 seeds, either dry (DS), after acquisition of secondary dormancy (0’) and after transfer on either 5mM KCl or KNO3 with and without ABA 50µM. MPK7 amount was monitored by immunoblot using an anti-MPK7 antibody. Equal loading was controlled by Coomassie staining of the membrane. LEAp, Late Embryogenesis Abundant proteins. Results were repeated two times depending on the time points.

### The MKK3 module does not modulate genes regulating ABA and GA contents

Last, we wondered whether the MKK3 module could modulate the ABA/GA balance. Therefore, we performed an RT-qPCR analysis of genes coding for ABA/GA biosynthetic and catabolic enzymes. These results are presented in figures S15 and S16. As expected, the expression of ABA synthesis genes (*NCEDs*, *ABA1*, *ABA2*, and *AAO3*) decreased after seed transfer to germination conditions, independently of NO_3_ . Genes involved in ABA catabolism had various patterns: *CYP707A1* and *CYP707A3* expression behaved like ABA biosynthetic genes; *CYP707A2* expression was expressed at least until 24 hours and, consistently with the literature (Yan et al., 2016), was strongly induced by NO_3_ ^-^; and *CYP707A4* expression was barely detectable in our conditions. We did not find any dramatic effect of the *mkk3* mutation on the expression of these ABA-related genes. GA biosynthetic genes *Ga20ox1/2/3* were neither NO_3_^-^- nor MKK3-dependent, whereas *Ga3ox* seemed to be promoted by NO_3_^-^- and *Ga3ox2* seems to present a delayed activation in *mkk3* (Figure S13).

## Conclusion and perspectives

Our work demonstrated that nitrate and light are activators of MKK3 module through the transcriptional regulation of two different pairs of MAP3K (figure 14). MAP3K13/14 function specifically in the regulation by nitrate whereas MAP3K19/20 seem rather to integrate both nitrate and light signals. We also showed that classical actors of dormancy regulation, such as NLP transcription factors and PHY photoreceptors, are involved in this activation. Last, we bring evidence that, in the context of secondary dormancy and with the assays used for this work, ABA and GA homeostasis are not the primary targets of the MKK3 pathway.

**Figure 13.**
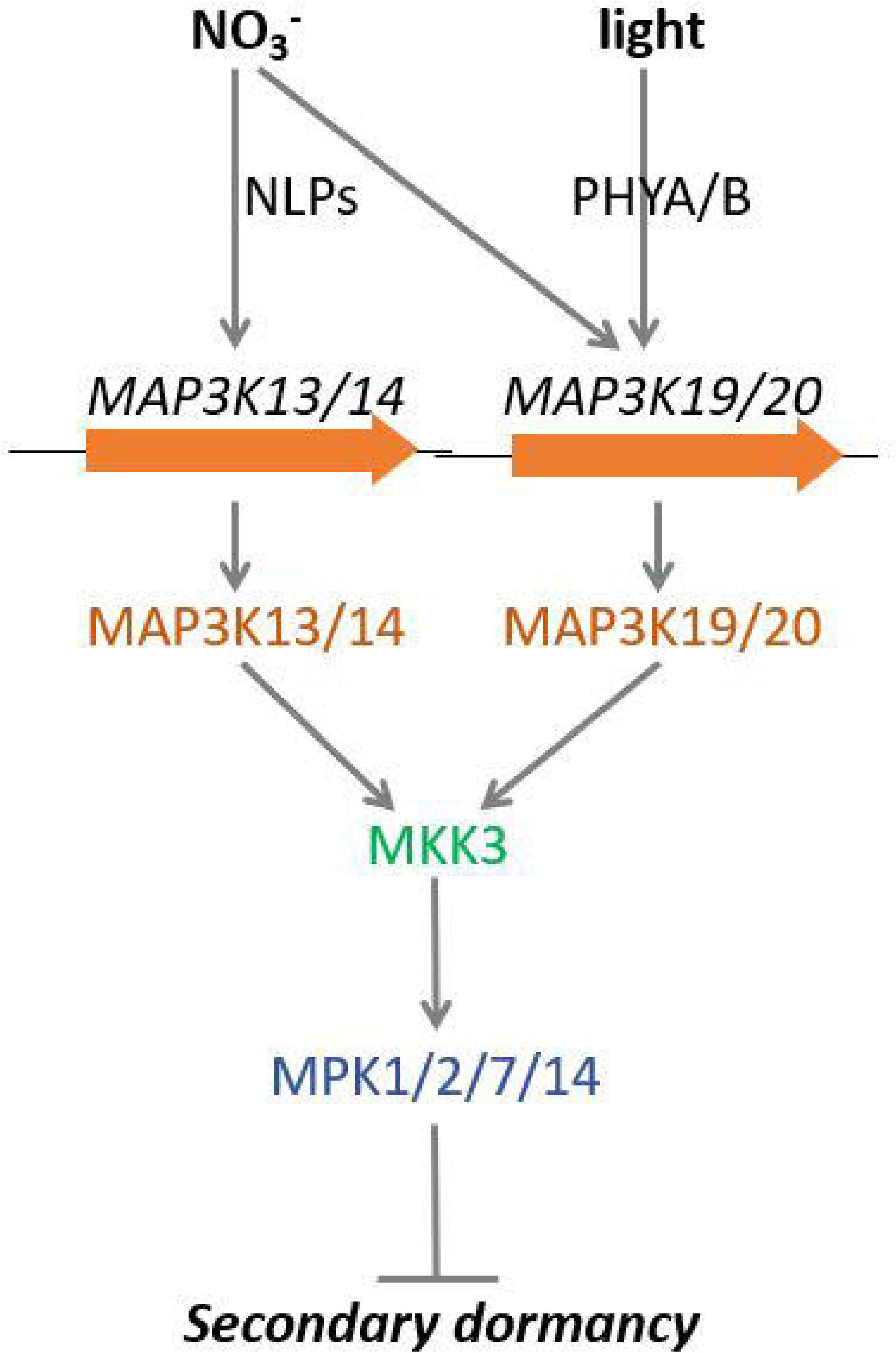
Working model of MKK3 module-dependent regulation of secondary seed dormancy by nitrate and light.

### Clade-III MAP3Ks, MKK3 and clade-C MAPKs emerge as a conserved transcription-dependent signalling module activated by a large range of signals

This work comforts the hypothesis that MKK3, together with clade-III MAP3Ks and clade-C MAPKs, defines robust signalling modules in plants (Colcombet et al., 2016). This specificity in kinase interaction is supported by yeast-2-hybrid and Split-YFP as well as functional reconstruction of modules in Arabidopsis protoplast expression system (Sözen et al., 2020; Danquah et al., 2015). Interestingly, the combined expression of MAP3K19, MKK3 and MPK7 in yeast strongly impaired cell growth, this impairment being suppressed if MAP3K19 or MPK7 is omitted or if MKK3 carries a mutation blocking its kinase activity (figure S17). This suggests that these three kinase clades are building a functional module in yeast and that MAP3K19 does not require further activation by a plant specific mechanism. The most striking demonstration that these kinases define functional modules comes from genetics and notably the observation that clade-C MAPK activities are impaired in mutants of upstream kinases. This impairment was complete in *mkk3* backgrounds whatever the activating signal used (Dóczi et al., 2007; Danquah et al., 2015; Sözen et al., 2020). In response to ABA and wound, clade-C MAPKs activities were also strongly reduced in mutants of clade-III MAP3Ks, *map3k17/18* for the first and *map3k14* for the second (Sözen et al., 2020; Danquah et al., 2015). In response to nitrate (in preparation) and nitrate/light (this study), the knocking out of all the transcriptionally-regulated MAP3Ks resulted in a total suppression of MAPK activation. This suggests once again that, at least for these signals, MAPK activation totally relies on clade-III MAP3KsOf course, these findings do not exclude the possibility MKK3 and clade-III MAP3Ks could also activate MAP2Ks/MAPKs of other sub-clades, as it has been suggested previously by other groups (Sethi et al., 2014; Benhamman et al., 2017; Lee, 2015; Bai and Matton, 2018; Takahashi et al., 2007; Schikora et al., 2008; Li et al., 2017a; Ojha et al., 2023; Takahashi et al., 2011). Notably MKK3 has been repeatedly to function upstream of MPK6 in response to light and pathogens (Takahashi et al., 2007; Schikora et al., 2008; Bai and Matton, 2018; Sethi et al., 2014; Lee, 2015) although co-expression in Arabidopsis protoplasts did not confirmed this connection (Danquah et al., 2015).

Our present work also shows that the module activation is systematically under the tight control of the signal-dependent transcriptional up-regulation of upstream MAP3K genes. In response to wound, systematically expression analysis identified 5 clade-III MAP3Ks, but when only a single one, *MAP3K14*, was knocked down, the mutant showed a MPK2 activity reduction (Sözen et al., 2020). We recently showed that nitrate transcriptionally regulates specifically *MAP3K13/14* genes in plantlets and that MPK7 activation was strongly impaired in the double mutant (Schenk, Chardin, Krapp, and Colcombet, unpublished). In the present work we generated a mutant in which each of the four *MAP3K* genes which are regulated in seeds transferred in germination-permissive conditions were mutated. In this quadruple mutant, the MPK7 activity is totally abolished in the same conditions. It is likely that the large number of clade-III MAP3Ks found in plant genomes (whereas there is usually a single MKK3 gene (Colcombet et al., 2016)) is evolutionarily constrained by the dual necessity of a large number of module activators and the importance of transcriptional regulation. Therefore, the MAP3K specificity should appear more at the promoter sequence level than at the protein sequence. Here we have identified light as a new signal able to activate the module. We showed that ABA, JA, nitrate and light are able to modulate the cascade but they are very likely other signals to be characterized. As a matter of fact, many more signals are able to regulate clade-III *MAP3K* expression as shown in Genvestigator expression database (Zimmermann et al., 2004; Colcombet et al., 2016) and some of the clade-III MAP3Ks have not described function yet.

### MKK3 module is a major dormancy regulator

The choice to germinate or not is crucial for seed survival. Understanding how plants make the decision, in interaction with their environment, has strong academic and agronomic interests. In this article, we report the dissection of a molecular signalling pathway integrating environmental cues to modulate secondary dormancy.

Our choice to work on secondary dormancy was methodological, but several arguments suggest that the MKK3 module may also function to release primary dormancy. (i) some clade-III *MAP3Ks* are strongly transcriptionally regulated during imbibition (Narsai et al., 2011), (ii) lines overexpressing some clade-III *MAP3Ks* and loss-of-function mutants display a germination phenotype (Choi et al., 2017; Mao et al., 2020; Danquah et al., 2015), and (iii) mutations in *MKK3* homologues have been identified in QTLs for wheat and barley vivipary, suggesting a direct mutation effect on primary dormancy (Torada et al., 2016; Nakamura et al., 2016). More directly, two recent articles support this idea. First, a whole MKK3 module plays an important role in the context of the temperature control of primary dormancy (Otani et al., 2024). Moreover a recent work in Arabidopsis reported that a functional MKK3-MPK7 module phosphorylate, in response to dormancy breaking conditions, the Ethylene Responsive Factor4 (ERF4) to target it to the proteasome. This allows the expression of EXPA genes necessary for the radicle emergence and seed germination. It is possible that the same players act downstream of our module described in the context of secondary dormancy (Chen et al., 2023).

Our work and others were led on the Brassicaceae specie *Arabidopsis thaliana* . It completes previous genetic investigations identifying *MKK3* mutation under important QTLs for barley and wheat pre-harvest sprouting (PHS) (Torada et al., 2016; Nakamura et al., 2016). These results on a dicotyledon suggest that the MKK3-dependent signalling pathway is shared among angiosperms and likely existed in the common ancestor of monocotyledons and dicotyledons.

Previously, we showed that ABA/drought and JA/wounding, through the production of MAP3K17/18 and MAP3K14, respectively, were activators of MKK3 and clade-C MAPKs (Colcombet et al., 2016; Sözen et al., 2020; Boudsocq et al., 2015). Our results also suggested that nitrate activates the module by upregulating *MAP3K13/14* (Schenk, Chardin, Krapp, and Colcombet, unpublished). Here, we unveiled that light can also activate the module and proposed that MAP3K19 and MAP3K20 could be the main entry points for this activation. A partial module composed of MKK3-MPK6 was shown to be activated by blue light to modulate MYC2-dependent photomorphogenesis (Sethi et al., 2014). Another study suggested, in the context of red light, that MKK3 rather restricts MPK6 activity in dark-light period transition (Lee, 2015). In the context of seeds, we also proposed that PHYA/B were the upstream light sensors, suggesting that the main compounds of the light effect are red and far-red wavelengths.

Besides clade-C MAPKs and MPK6, MKK3 has been proposed to work upstream of a clade-D MAPK, MPK8, in the context of wound-triggered Reactive Oxygen Species (ROS) homeostasis (Takahashi et al., 2011). Interestingly, a recent study also highlighted the role of MPK8 in regulating seed dormancy (Zhang et al., 2019). In this work, the authors show that freshly harvested *mpk8* seeds display a strong dormancy phenotype, arduously released by gibberellins and after-ripening. MPK8 can interact with and phosphorylate the transcription factor TEOSINTE BRANCHED1/CYCLOIDEA/PROLIFERATING CELL FACTOR (TCP14) in the nucleus, enhancing the activity of the latter in seeds. Transcriptomes of WT, *mpk8*, and *tcp14* seeds are very similar at the dry stage but diverge after a 24 h imbibition. Nevertheless, *mpk8* and *tcp14* mutants displayed a strong overlap in their misregulated genes, confirming that the two proteins belong to a common signalling pathway. The connection between MKK3 and MPK8 has not been confirmed yet, since MKK3 does not reproducibly activate MPK8 in protoplasts (Danquah et al., 2015). Nonetheless, identifying how MPK8 and TCP14 are regulated in our conditions would be interesting

Surprisingly, ABA is not an activator of the module in seeds, whereas it has been repeatedly reported to be in plantlets and adult plants (Mitula et al., 2015; Matsuoka et al., 2015; Danquah et al., 2015). We observed neither an upregulation of *MAP3K17/18* or other clade-III *MAP3Ks*, nor the increase of MPK7 activity in dormant seeds transferred to germination conditions supplemented with ABA. This result suggests that MAP3K promoters are specifically recognised by ABA-responsive transcription factors, which are not recruited by ABA in seeds or not expressed. Because ABA and nitrate/light act in opposite ways to promote seed germination, it makes sense that both signals cannot activate the same module. We also tested the activation by GA or its importance in MPK7 activation by light/nitrate but did not observe any effects of GA or paclobutrazol. This suggests that the module integrates environmental signals to modulate cellular mechanisms controlling germination. One possibility is that it modulates the ABA/GA balance by regulating ABA/GA biosynthetic genes.

## Material and methods

### Biological material

*mkk3-1* (SALK_051970), *mkk3-2* (SALK_208528), *mpk7-1* (SALK_113631) and lines expressing HA-tagged *MPK1/2/7* loci were published previously (Dóczi et al., 2007; Sözen et al., 2020). *mpk1-1/2-2/7-1/14-1* and *map3k19-1/20-3* were obtained by crossing in the Pr Kawakami’s laboratory from the following single mutant lines *mpk1-1* (SALK_063847C)*, mpk2-2* (SALK_047422C)*, mpk7-1* (SALK_113631)*, mpk14-1* (SALK_022928C), *map3k19* (Transposon pst14411)*, map3k20-3* (CS443915/GK-458D07) (Otani et al., 2024)*. map3k13CR/14CR* are a double crisper mutants described previously (Sözen et al 2020; Schenk, Chardin, Krapp, and Colcombet, unpublished). *map3K13CR*/*14CR/19-1/20-3* were obtained by crossings. *nlp8* (SALK_16341), *nlp7/nlp8* (SALK_026134/SALK_16341) and nlp8/nlp9 (SALK_140298/SALK_025839) were published previously (Yan et al., 2016). *nlp7/nlp8/nlp9* (SALK_026134/SALK_031064/SALK_025839) was created by crossing.

To produce *35S::MAP3K19-MYC* lines, the ORF was PCR amplified from Arabidopsis thaliana (ecotype Columbia-0) cDNA using iProof DNA polymerase (Bio-Rad), specific primers (ORF-MAP3K19-F: gga gat aga acc ATG GAG TGG ATT CGA GGA GAA A; ORF-MAP3K19-R tcc acc tcc gga tcm CCG TAC GGT GAC CCA GCT) and a two-step amplification protocol as described previously (Colcombet et al., 2013). PCR products were recombined into pDONR207 (Invitrogen) using Gateway® BP Clonase® II Enzyme mix (Invitrogen). LR recombination reactions were performed using Gateway® LR Clonase® Enzyme Mix (Invitrogen) in order to transfer ORF sequences from Entry vectors to the pC2N1 allowing C-terminal translational fusion with the 10xMyc tag under the control of the 35S promoter (Bigeard et al., 2014; Berriri et al., 2012). The resulting construct, *pC2N1-MAP3K19,* was introduced into the *Agrobacterium tumefaciens* strain C58C1 and used to transform *Arabidopsis thaliana* Col-0 plants by the floral-dipping method (Clough and Bent, 1998). Using kanamycin segregation analysis, we selected two independent transgenic lines carrying a single insertion at the homozygous state. *MAP3K19-MYC* expression was assessed by western blot.

### Growth production

Seeds were produced in ‘low-nitrogen’ conditions (Alboresi et al., 2005). They were germinated on agar media on ½ MS plates in a growth chamber in long-day conditions for 7 days, and plantlets were transferred onto ‘Spezialsubstrat’ (Stender, ref: 19002774-A204 sans NPK) containing low nitrate. Plants were further grown in an Aralab® growth cabinet maintained in long-day conditions (16 h of 80–100 μE m^-2^ s^-1^ light at 20°C and 8 h of darkness at 18°C) with a 60% hygrometry. Pots were watered three times weekly (Monday, Wednesday, and Friday), using a low-nitrogen solution (250 µM KH_2_PO_4_; 250 µM MgSO_4_; 750 µM KNO_3_; 125 µM Ca(NO_3_)_2_; 125 µM CaCl_2_; 10 mG L^−1^ Sequestrene138FE 100SG (Syngenta); 0,4 µM (NH_4_)_6_Mo_7_O_24_; 243 µM H_3_BO_3_; 118 µM MnSO_4_; 34,8 µM ZnSO_4_; 10 µM CuSO_4_) containing 1 mM NO_3_^−^ (Loudet et al., 2003) and filling the plateau with 2 cm of solution. After 1–2 hours, the remaining liquid was removed by draining. In these growth conditions, plantlets exhibit smaller rosettes and complete their lifecycle after 3–4 months. When about two-thirds of the siliques started turning yellow, watering was stopped, and plants dried. The inflorescence was cut, enclosed in paper bags, and further dried for two weeks. Seeds were then harvested and stored at room temperature in Eppendorf tubes.

### Dormancy induction and germination tests

Appropriate amounts of seeds were sterilised (15’ in [50% EtOH, 0.5% Triton 100X], 2x 5’ in 96% EtOH). Seeds were dried on sterile Whatman paper (ref: GE Healthcare Life Science, ME 25/31 ST, Whatman). 5-cm round plates were filled with 8 mL minimal medium (MES hydrate [M8250, Sigma Aldrich] 0.58%, Agar [HP696, Kalys 0,7%], pH adjusted to 5,75 with NaOH). Depending on the type of experiment performed, the minimal medium was supplemented with KNO_3_, KCl, 50 µM ABA (Sigma Aldrich, ref: A1049, in ethanol), 50 µM GA_3_ (Sigma Aldrich, ref: G7645), or 10 µM Paclobutrazol (Sigma Aldrich, ref: 43900) at indicated concentrations. A round filter (GE Healthcare Life Science ME 25/31 ST) was carefully placed on the medium surface for experiments requiring seed transfer. Typically, 50–150 seeds were sown on each plate. Plates were sealed with micropore surgical tape.

To induce secondary dormancy, plates were wrapped in aluminium foil and incubated at 30°C in a Memmert cabinet for 1 to 15 days. To evaluate dormancy through the ability of seeds to germinate, plates were shifted in a growth room in long-day conditions (16 h light [80–100μE m^-2^s^-1^] at 22°C and 8 h dark at 18°C). To evaluate the ability of anions or hormones to release dormancy, dormant seeds were delicately transferred onto new media with appropriate supplementation. The germination rate was expressed as the percentage of seeds with a radicle protrusion after seven days.

### Gene expression

For gene expression analysis, seeds were collected, frozen in liquid nitrogen, and ground using a plastic pestle. RNA was extracted using the NucleoSpin® RNA Plant kit (Macherey-Nagel) according to the manufacturer’s instructions and quantified with a Nanodrop spectrophotometer. Typically, 2–5 µg of total RNA was used to perform RT, using the Transcriptase inverse SuperScript™ II (Thermofisher) and following the manufacturer’s instructions. 10 ng of cDNA was used for qPCR with the CFX384 Touch real-time PCR detection system (Bio-Rad) and ONEGreen® Fast qPCR Premix (Ozyme), following the manufacturer’s standard instructions. *ACT2 (*AT3G18780) was used as an internal reference gene to calculate relative expression. RT-qPCR primers are listed in Supplemental Table 1.

### Kinase assay and western blot

For biochemistry, 50–150 seeds were collected, frozen in liquid nitrogen, and ground using a plastic pestle. A detailed kinase assay protocol for plant samples has been provided in a previous publication (Sözen et al., 2020). The notable modification necessary for protein extraction from seeds is that triton was adjusted at 1% in the extraction buffer. A new batch of rabbit α-MPK7 antibodies was also prepared for this study, raised against the LYYHPEAEISNA epitope. For kinase assays, immunoprecipitated kinases were resuspended in 15 µl of kinase buffer containing 0.1 mM ATP, 1 mg ml^-1^ MBP, and 2 µCi ATP [γ-33P]. After 30 min of reaction at room temperature, retain was stopped with 15µL Laemmli buffer 2x and boiled for 5 minutes at 95°C. Samples were then loaded and run on a 15% SDS-PAGE gel. MBP was stained with Coomassie Blue. Then, the gels were dried and revealed on a STORM scanner (GE Healthcare).

Protein levels were monitored by immunoblotting following Bio-Rad recommendations. Proteins were separated in 10% (w/v) SDS-PAGE gels and transferred onto polyvinylidene fluoride membranes (Bio-Rad). Membranes were blocked in 5% (w/v) nonfat dry milk. The following primary and secondary antibodies were used: α-HA (Roche 11867431001; 1: 10,000 dilution), α-Myc (Sigma-Aldrich C3956; 1:10,000 dilution), α -rat (Sigma-Aldrich A9037, 1:10,000), α-rabbit (Sigma-Aldrich A6154, 1/20,000), and α -mouse (Sigma-Aldrich A5906, 1:10,000). Horseradish peroxidase activity was detected with a Clarity western ECL substrate reaction kit (Bio-Rad) and a ChemiDoc Imagers (Bio-Rad). Blots were stained with Coomassie Blue for protein visualisation.

### Functional expression of kinase in yeast

*MAP3K19*, *MKK3* and *MPK7* ORFs were amplified using couple of primers and cloned into p425GPD, p426GPD and p423GPD (Mumberg et al., 1995)respectively. To generate MKK3 mutants, site-directed mutagenesis was carried out using the QuickChange Lightning kit from Agilent. The two targeted mutation sites were K112M and D207A. Combination of plasmids were co-transformed in B4741 *Y00000* (*MAT a; his3D1; leu2D0; met15D0; ura3D0* ) strain derived from the S288C isogenic yeast strain. Transformed yeasts were selected and grown on selective medium (Yeast Nitrogen Base, 2 % glucose, and the addition of amino acids except histidine, uracil and leucine to maintain selection for the plasmids).

## Acknowledgements

We thank our fellow scientists from the “Stress Signalling” team and more broadly from our research laboratories for constructive scientific discussions. We also thank from the bottom of our hearts our colleagues who are not referred in the author list above but brought us the precious technical, logistical and administrative support necessary for an effective research activity. Viviane Bréhaut is thanked for her help in building mutants and Eiji Nambara for *nlp* seeds. This work was directly supported by the French State through the ANR MAPKSEED (ANR-18-CE20-0019) and the LabEx Saclay Plant Sciences-SPS (ANR-10-LABX-0040-SPS). Cécile Sözen is thanked for the critical reading/editing of the manuscript.

## Figure legends

**Figure S1 - Seeds impaired in MKK3 have a reduced nitrate-triggered dormancy release**

A and B. Seeds from indicated genetic background were imbibed at 30°C in the dark for 10 days to induce secondary dormancy and transferred on medium containing indicated NO3 concentration. Germination ability was assessed after 7 days in long day conditions. Values are mean ± SE of two biological replicates from seed batches produced independently. Values for each replicate are also shown. On the top, based on Mann-Whitney test, yellow sticks show differences with α < 5% whereas gray sticks show no differences.

**Figure S2 - MPK7 is expressed in dry seeds**

Western-Blot detection in dry seeds and 7-days old plantlets of lines expressing indicated HA tagged clade-C MAPK from the native locus. Equal loading was controlled by Coomassie staining of the membrane. LEAp, Late Embryogenesis Abundant proteins. RbcL, RubisCo large subunit. Results were repeated twice.

**Figure S3 - Quantification of MPK7 activity in dormant Col-0 and *mkk3-1* seeds transferred on nitrate.**

MPK7 activities of replicates were quantified and normalized to the one of Col-0 8h NO3 (*). Values are mean ± SE of two to six biological replicates from seed batches produced independently. Values for each replicate are also shown.

**Figure S4 - kinase activity immunoprecipitated with the anti-MPK7 antibody dependent of the expression of MPK7.**

Kinase activity of MPK7 after immunoprecipitation with an anti-MPK7 antibody from Col-0 and *mpk7-1* seeds, either dry (DS), after acquisition of secondary dormancy (0’) and after transfer on either 5mM KCl or KNO3. Seeds were produced in the green house with non limiting nitrogen fertilizer, probably explaining why there is no NO3 effect.

**Figure S5 - MPK7-HA activity in dormant seeds expressing an HA-tagged MPK7 depends on MKK3.**

Kinase activity of MPK7 after immunoprecipitation with an anti-HA antibody from *MPK7locus-HA* and *mkk3-1 MPK7locus-HA* seeds, either dry (DS), after acquisition of secondary dormancy (0’) and after transfer on either 5mM KCl or KNO3. MPK7-HA amount was monitored by immunoblot using an anti-HA antibody. Equal loading was controlled by Coomassie staining of the membrane. LEAp, Late Embryogenesis Abundant proteins.

**Figure S6 - All clade-III MAP3Ks are not expressed in seeds or during secondary dormancy release**

RT-qPCR analysis of *MAP3K15, MAP3K16* , *MAP3K17*, *MAP3K18* and *MAP3K21* genes expression. Transcript levels are expressed relative to ACTIN2 as reference gene. Values are mean ± SE of two to 10 biological replicates from seed batches produced independently. Values for each replicate are also shown.

**Figure S7 - Quantification of MPK7 activity in dormant Col-0 and map3k mutant seeds transferred on nitrate.**

MPK7 activities of replicates were quantified and normalized to the one of Col-0 8h NO3 (*). Values are mean ± SE of 2-6 (A) or 3 (B) biological replicates from seed batches mainly produced independently. Values for each replicate are also shown.

**Figure S8 - Quantification of MPK7 activity in Col-0 and *35S::MAP3K19-MYC* seeds transferred on nitrate.**

MPK7 activities of replicates were quantified and normalized to the one of Col-0 8h NO3 (*). Values are mean ± SE of one to three biological replicates from seed batches produced independently. Values for each replicate are also shown.

**Figure S9 – Quantification of MPK7 activity in dormant Col-0 and *nlp8* seeds transferred on nitrate.**

MPK7 activities of replicates were quantified and normalized to the one of Col-0 8h NO3 (*). Values are mean ± SE of two to five biological replicates from seed batches produced independently. Values for each replicate are also shown.

**Figure S10 - Quantification of MPK7 activity in Col-0 and *nlp* seeds transferred on nitrate.** MPK7 activities of replicates were quantified and normalized to the one of Col-0 8h NO3 (*). Values are mean ± SE of 3 biological replicates from seed batches produced independently. Values for each replicate are also shown.

**Figure S11 - Quantification of MPK7 activity in Col-0 transferred on nitrate and dark.**

MPK7 activities of replicates were quantified and normalized to the one of Col-0 8h NO3 (*). Values are mean ± SE of three to five biological replicates from seed batches produced independently. Values for each replicate are also shown.

**Figure S12 – Complement of clade-III *MAP3K* expression in seeds or during secondary dormancy release by nitrate and/or light**

RT-qPCR analysis of *MAP3K15/16/17/18/21* genes expression. Transcript levels are expressed relative to ACTIN2 as reference gene. Values are mean ± SE of three biological replicates from seed batches produced independently. Values for each replicate are also shown. ND, not determined.

**Figure S13 - Quantification of MPK7 activity in Col-0 seeds after transfer on Paclobutrazol (PCZ) and GA3.**

A and B. MPK7 activities of replicates were quantified and normalized to the one of Col-0 8h NO3 (*). Values are mean ± SE of two biological replicates from seed batches produced independently. Values for each replicate are also shown.

**Figure S14 – Expression of *MAP3K13*, *MAP3K14*, *MAP3K17, MAP3K18, MAP3K19* and *MAP3K20* is not affected by ABA during secondary dormancy release**

RT-qPCR analysis of *MAP3K13*, *MAP3K14*, *MAP3K17*, *MAP3K18*, *MAP3K19* and *MAP3K20* genes expression. Transcript levels are expressed relative to ACTIN2 as reference gene. Values are mean ± SE of two biological replicates from seed batches produced independently. Values for each replicate are also shown.

**Figure S15 – Expression of genes involved in ABA biosynthesis (left) and catabolism (right)** RT-qPCR analysis of indicated genes expression in Col0 and *mkk3-1*. Transcript levels are expressed relative to ACTIN2 as reference gene. Values are mean ± SE of two to three biological replicates from seed batches produced independently. Values for each replicate are also shown.

**Figure S16 – Expression of genes involved in ABA metabolism**

RT-qPCR analysis of indicated genes expression in Col0 and *mkk3-1*. Transcript levels are expressed relative to ACTIN2 as reference gene. Values are mean ± SE of two to three biological replicates from seed batches produced independently. Values for each replicate are also shown.

**Figure S17 – reconstruction of a functional MKK3 pathway in budding yeast strongly affects its growth.**

Serial dilutions of the various transformed strains were spotted onto agar-based solid medium. Note that the complete MKK3 module leads to growth inhibition. However, the growth is restored if MAP3K19 or MPK7 is omitted, or if MKK3 carries a kinase-dead mutation affecting its kinase activity.

---

**Supplemental Table 1.**
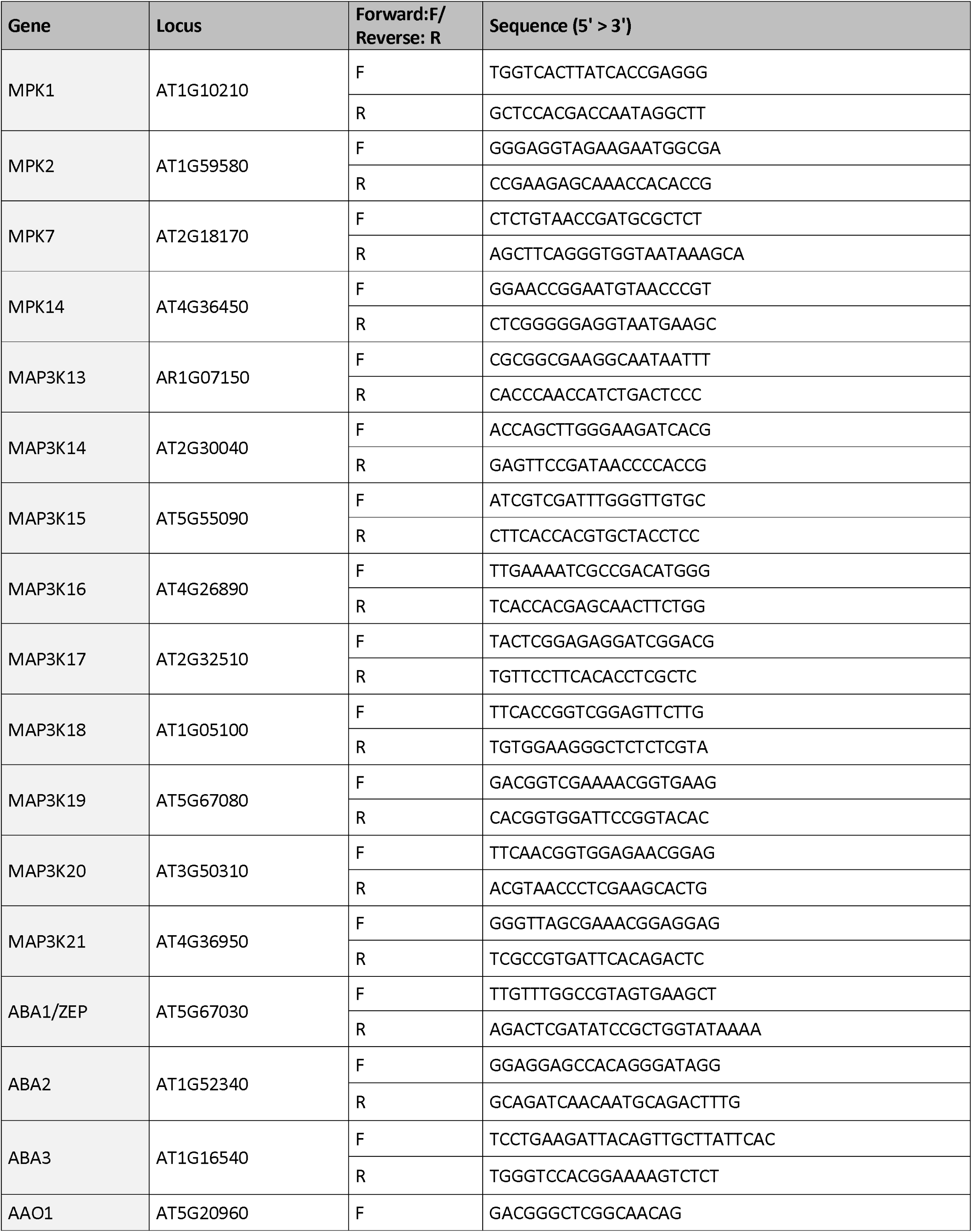
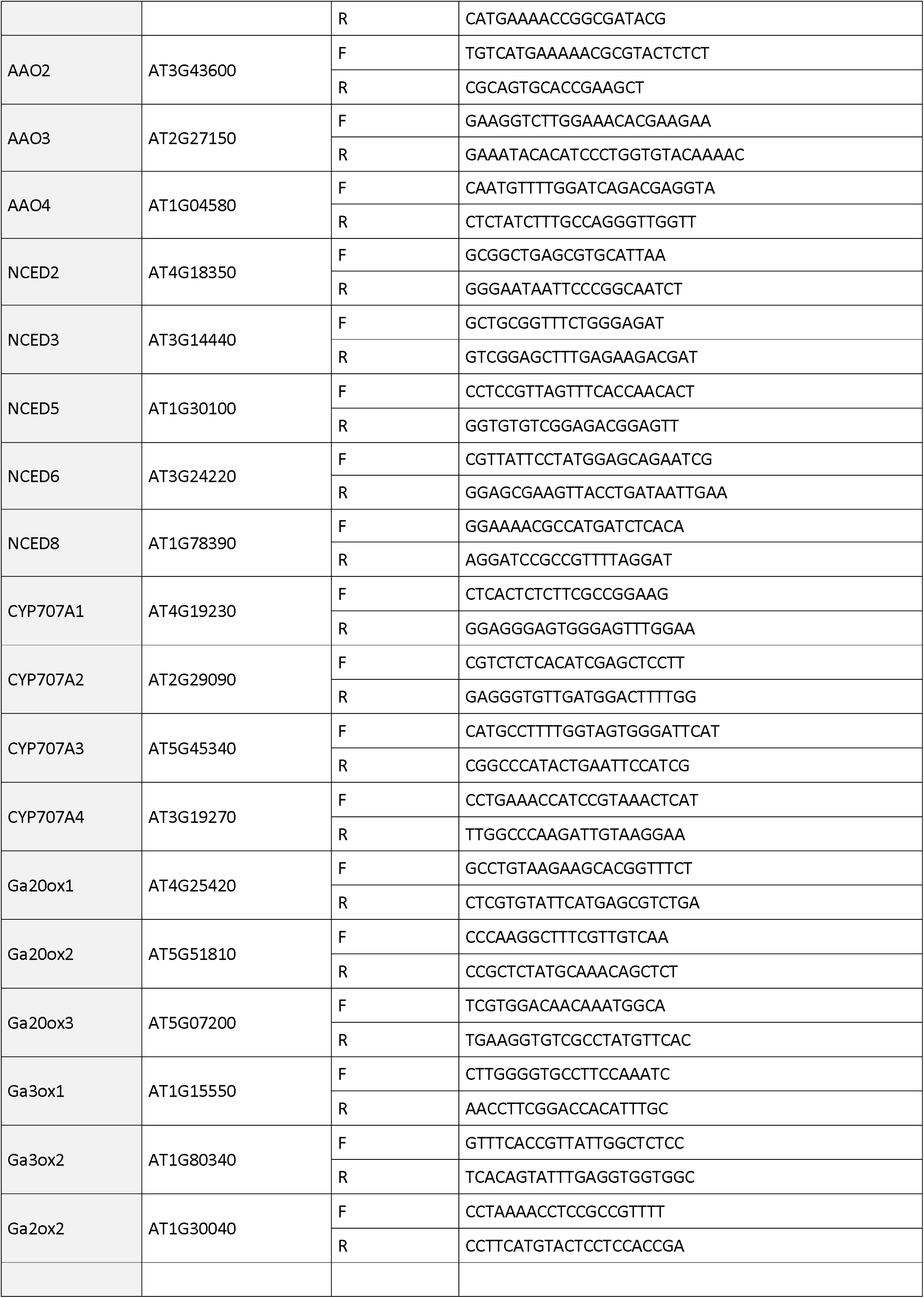
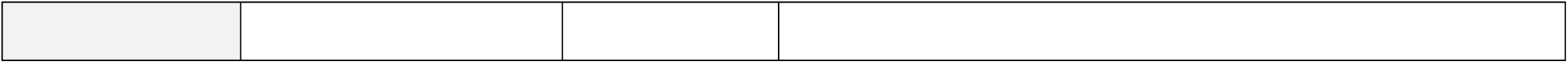

